# Primary tuberculous mycobacterial granulomas provide a niche for superinfecting *Mycobacterium abscessus*

**DOI:** 10.1101/2025.05.05.652332

**Authors:** Denise Wee, Manitosh Pandey, Yao Chen, Paolo A Lorenzini, Eve WL Chow, Yue Wang, Amit Singhal, Stefan H Oehlers

## Abstract

Prior and concurrent tuberculosis infection are among the most important susceptibility factors for nontuberculous mycobacterial infection in Asia. Here we model this process in zebrafish with a primary *Mycobacterium marinum* infection followed by a secondary *M. abscessus* infection. We demonstrate preferential growth of secondary *M. abscessus* infection inside primary *M. marinum* granulomas. Granuloma-resident secondary *M. abscessus* is protected from macrophage-mediated immune control and antibiotic therapy. Successful colonization is driven by expansion of *M. abscessus* feeding on caseum produced by the primary *M. marinum* ESX-1 virulence program in a nutritionally separate niche from *M. marinum*. Our data suggest tuberculous granulomas may provide a long-lasting niche for the growth of the opportunistic pathogen *Mycobacterium abscessus*.

## Introduction

Infections by *Mycobacterium abscessus* are an emerging global health problem, and the rising rates of infection by similar non-tuberculous mycobacteria (NTM) threaten to undermine the progress made in tuberculosis (TB) control in many countries (*1*). Post-TB lung disease is an important but understudied chronic respiratory disease that contributes to mortality and morbidity, including raising the risk of subsequent NTM and TB infections in successfully treated TB patients (*2, 3*). In countries with endemic TB, NTM-TB coinfection accounts for up to 10% of the NTM patient population (Shandong Province, China) and 3% of the TB patient population (Taiwan) (*4, 5*). This association contrasts with the known protective effect of homologous *M. tuberculosis* infection against *M. tuberculosis* superinfection and the heterologous protection conferred by the *M. bovis* BCG vaccine against subsequent mycobacterial infections.

The granuloma is the histological hallmark of TB infection and serves as the primary immunological interface between pathogenic mycobacteria and the host immune system. These long-lasting structures persist despite immune and antibiotic containment of *M. tuberculosis*, with varying estimates of their role in maintaining a reservoir for subclinical TB. In the case of calcified granulomas, they can remain lifelong (*6, 7*). Evidence that granulomas can harbor superinfecting tuberculous mycobacteria spans both experimental and clinical datasets, including key milestone studies demonstrating homing of superinfecting *Mycobacterium marinum* into zebrafish and frog granulomas, *M. tuberculosis* and *M. bovis* into granulomas in mice and humans, respectively, and a high rate of cutaneous granulomas containing multiple mycobacterial species (*8-12*). Intriguingly, NTM have been detected in IS*6110* positive TB granuloma biopsy tissues, demonstrating infiltration of NTM into *M. tuberculosis* granulomas (*13*). However, the role of tuberculous granulomas in driving susceptibility to NTM infection has yet to be examined. We hypothesized organized caseated granulomas may act as a protective niche for superinfecting mycobacteria, shielding them from cross-protective immune control and providing a readily available source of nutrients for rapid NTM growth.

Here, we demonstrate that colonization with the natural mycobacterial pathogen *M. marinum* increases the susceptibility of zebrafish to *M. abscessus*. This susceptibility can be directly attributed to macrophage-mediated, ESX1-dependent carriage of *M. abscessus* into pre-existing caseous granulomas, where it is shielded from containment by the host immune system. We find that *M. abscessus* adapts to growth in the necrotic granuloma, enabling occupancy of a distinct nutritional niche.

## Results

### Primary *M. marinum* infection predisposes adult zebrafish to secondary *M. abscessus* infection

Previous studies using the frog-*M. marinum* and macaque-*M. tuberculosis* models have demonstrated protection against homologous superinfection (*8, 14, 15*). The adult zebrafish platform provides a unique intersection of tuberculous granuloma formation when challenged with *M. marinum* and an immunologically-intact permissive host for *M. abscessus* (*16, 17*). We performed a sequential infection experiment in adult zebrafish to determine if primary tuberculous infection promotes resistance or susceptibility to secondary *M. abscessus* infection. A high-dose secondary infection with of 10^6^ CFU of *M. abscessus* caused unexpected mortality in animals with a primary *M. marinum* infection (Figure 1A). Animals infected with a primary *M. marinum* infection and a lower-dose secondary infection with 5x10^5^ CFU of *M. abscessus* exhibited a higher *M. abscessus* burden than naïve animals (Figure 1B).

**Figure 1.**
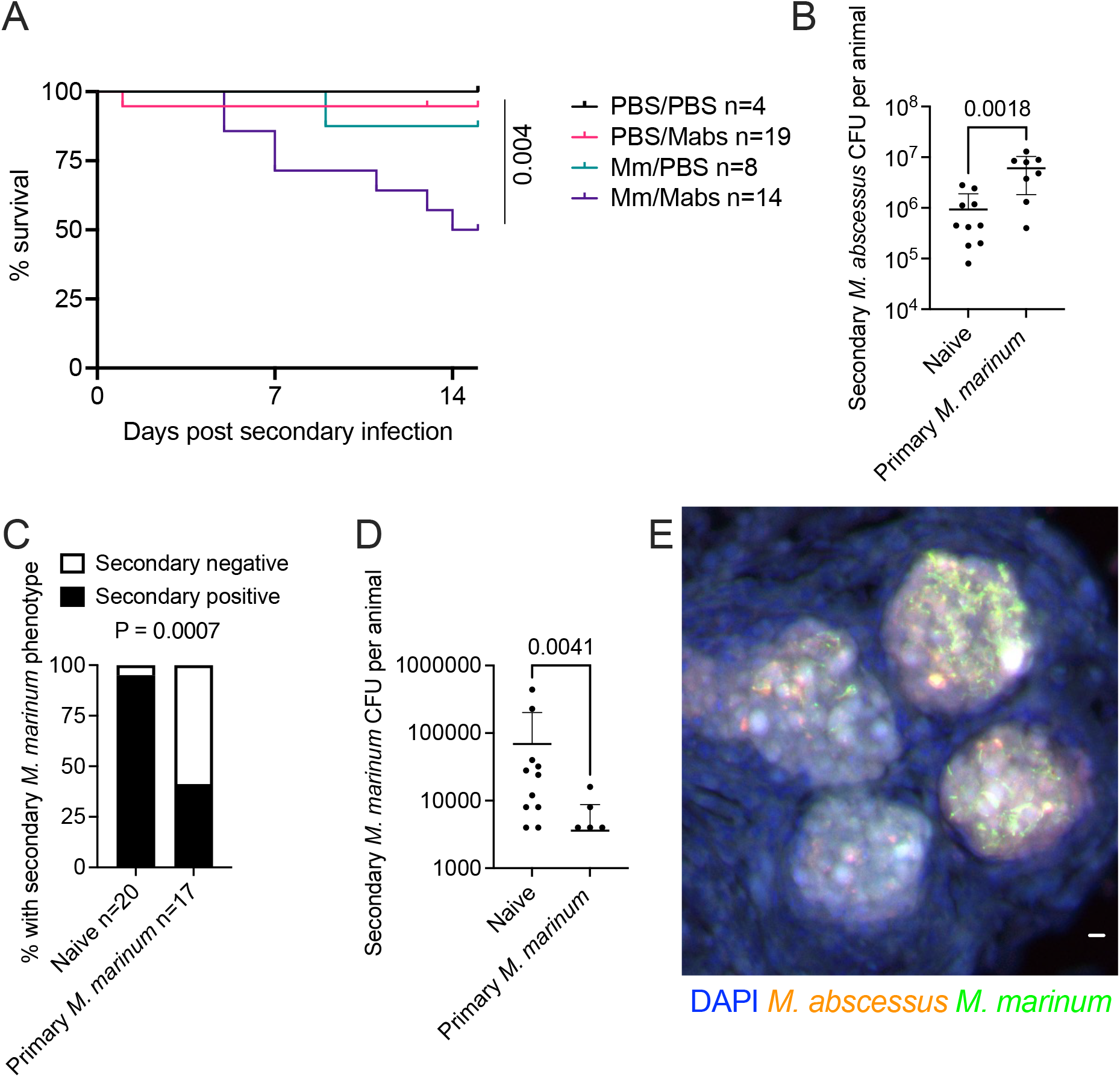
Pre-existing *M. marinum* infection worsens secondary *M. abscessus* infection but protects from secondary *M. marinum* infection A.Survival curve of adult zebrafish infected with combinations of primary *M. marinum* (Mm) and secondary *M. abscessus* (Mabs) at 2 weeks post primary infection. Data is from a single experiment with 45 animals. Statistical analysis reported is a Log-rank test comparing the PBS/Mabs group with the Mm/Mabs group. B.Quantification of secondary *M. abscessus* burden at 2 weeks post secondary infection in naïve and primary *M. marinum-*infected adult zebrafish infected at 2 weeks post primary infection. C.Quantification of secondary *M. marinum* engraftment at 2 weeks post secondary infection in naïve and primary *M. marinum-*infected adult zebrafish infected at 2 weeks post primary infection. Data is pooled from two biological replicates. Statistical analysis reported is a Fisher’s Exact Test. D.Quantification of secondary *M. marinum* burden at 2 weeks post secondary infection in naïve and primary *M. marinum-*infected adult zebrafish infected at 2 weeks post primary infection. E.Representative image of multilobed granuloma in 2 week post secondary infection adult zebrafish infected with secondary *M. abscessus* at 2 weeks post primary *M. marinum* infection. Scale bar indicates 10 μm.

Mammalian models of *M. tuberculosis* sequential infections have demonstrated significant protection from infection (*8, 14, 15, 18*). We performed similar homotypic experiments with sequential *M. marinum* infection to determine if our *M. abscessus* phenotype was an artifact of an exhausted immune system. We found that primary *M. marinum* infection protected against a homotypic secondary *M. marinum* infection both by preventing the engraftment of a low-dose *M. marinum* infection (Figure 1C) and reducing the recovered *M. marinum* burden from a higher dose infection (Figure 1D).

Cryosectioning of sequentially infected fish was performed to determine the spatial localization of superinfecting *M. abscessus* relative to primary *M. marinum* granulomas. Consistent with the homologous superinfection literature (*8*), *M. marinum* granulomas were found to be colonized by secondary *M. abscessus* (Figure 1E).

### Colonization of primary *M. marinum* granulomas accelerates the growth of superinfecting *M. abscessus*

While the adult zebrafish experimental system facilitates the study of classic hypoxic caseous necrotic granulomas (*19-21*), it does not readily facilitate real-time imaging of host-microbe interactions, as can be achieved with zebrafish embryos. Furthermore, sequential intraperitoneal injections likely bath the primary granuloma in secondary *M. abscessus* facilitating ready uptake. For these reasons we switched to the use of the zebrafish embryo-*M. marinum*/*M. abscessus* infection system to study the mechanisms of primary *M. marinum* infection-mediated susceptibility to secondary *M. abscessus* infection. We established an experimental sequential infection system in zebrafish embryos by performing the primary injection into the neural tube at the previously characterized “trunk” injection site (*21*), then performing secondary injection into the circulation at 3 days post primary infection (dppi) (Figure 2A). Timelapse imaging revealed uptake of *M. abscessus* from the circulation into established *M. marinum* granulomas (Figure 2B and Supplementary Video 1).

**Figure 2.**
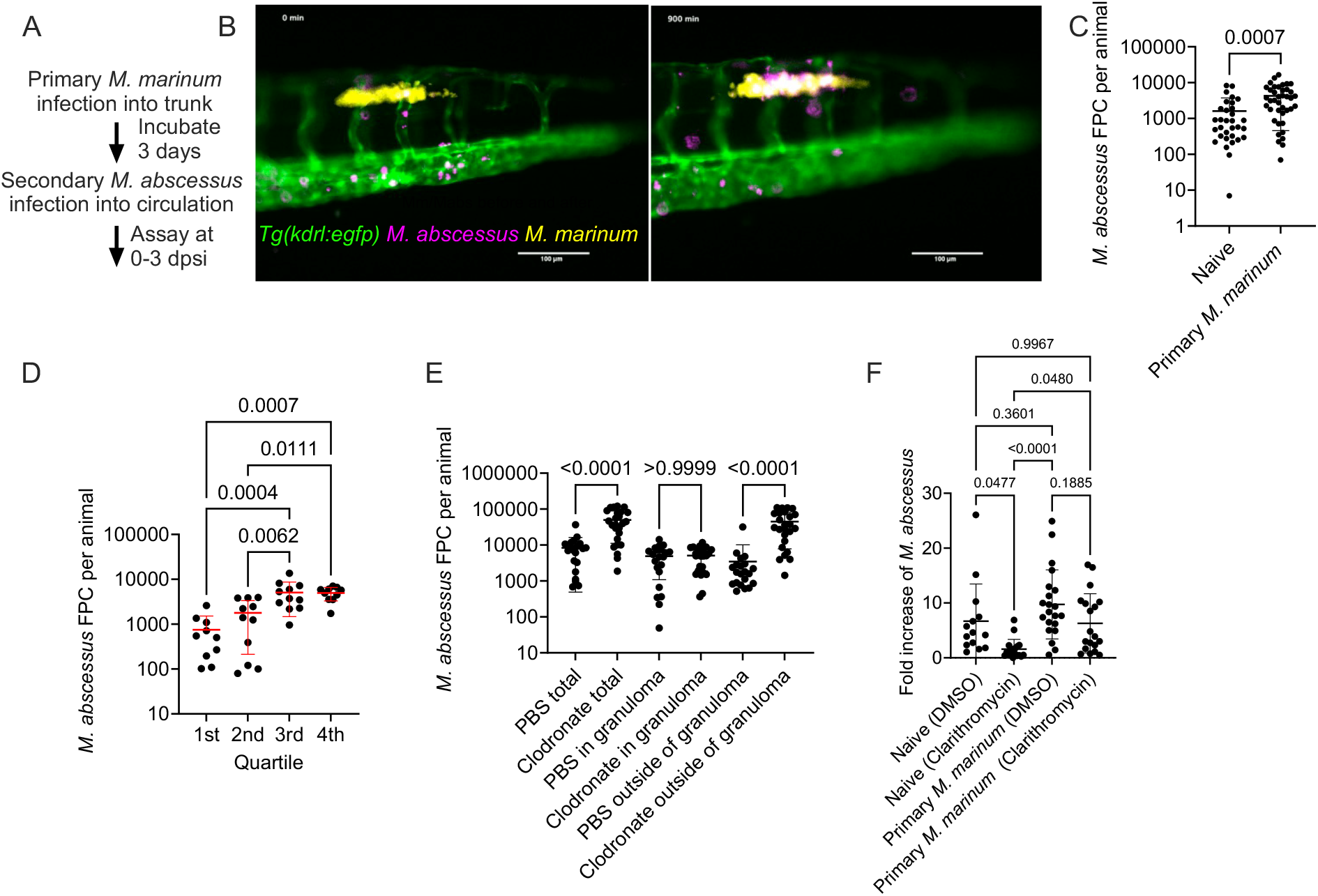
Ingress into *M. marinum* granulomas protects *M. abscessus* from innate immunity A.Schematic illustrating embryo superinfection experimental set up. Abbreviation: Days post secondary infection (dpsi). B.Still images extracted 900 minutes apart from Supplementary Video 1. Site of *M. abscessus* (purple) injection is indicated within *Tg(kdrl:egfp)-*positive vasculature (green) below primary *M. marinum* granuloma (yellow). Scale bars indicate 100 μm. C. Quantification of secondary *M. abscessus* burden in 3 dpsi embryos infected with primary *M. marinum*. D. Quantification of secondary *M. abscessus* burden in 3 dpsi embryos infected with primary *M. marinum* grouped by quartile of *M. abscessus* found within *M. marinum* granulomas. E. Quantification of secondary *M. abscessus* burden in 3 dpsi embryos infected with primary *M. marinum, M. abscessus* was delivered by co-injection with clodronate microsomes. F. Quantification of fold change in *M. abscessus* burden during treatment with 10 μM clarithromycin from 2 dpsi to 4 dpsi.

Consistent with our adult zebrafish data, fluorescent pixel count revealed an increased *M. abscessus* burden in embryos 3 days post secondary infection (dpsi), when sequentially infected at 3 dppi with *M. marinum*, compared to *M. abscessus* infection of naïve embryos (Figure 2C). Furthermore, analysis of *M. abscessus* burden in sequentially infected embryos grouped by quartile of colonization revealed a higher overall *M. abscessus* burden in embryos with a high rate of colonization (Figure 2D). Together, these data suggested colonization of primary granulomas accelerated secondary *M. abscessus* growth.

To examine the contribution of prior *M. marinum* granuloma formation relative to more general *M. marinum*-mediated immune subversion, we compared *M. abscessus* growth following mono-and co-injection with *M. marinum*. Unlike our sequential injection infection data, co-injection of *M. marinum* did not affect the growth of *M. abscessus* in zebrafish embryos, demonstrating the importance of granuloma formation in driving susceptibility to sequential *M. abscessus* infection (Supplementary Figure 1A). Interestingly, we found a contrasting antagonistic effect of *M. abscessus* infection on *M. marinum* growth, reminiscent of a trained immunity effect (Supplementary Figure 1B).

To determine if immune pressure restricts *M. abscessus* growth, resulting in preferential growth within primary *M. marinum* granulomas, we co-injected clodronate with secondary *M. abscessus* to deplete macrophages and dexamethasone immune suppression to alleviate immune pressure. Total *M. abscessus* burden increased with clodronate co-injection, driven entirely by an increase in the extra-granuloma compartment. This finding is consistent with our hypothesis that granulomas provide *M. abscessus* with a physical sanctuary from the host immune control (Figure 2F).

Caseum increases mycobacterial antibiotic tolerance by excluding antibiotics and rewiring bacterial physiology (*22, 23*). Specifically, stationary phase *M. abscessus* in caseum is markedly more resistant to frontline antibiotics such as bedaquiline, clarithromycin, imipenem, clofazimine, and moxifloxacin compared to actively growing broth cultures (*23*).. Clarithromycin, an example of a current front line antibiotic, was effective at restraining the growth of *M. abscessus* in naïve but not *M. marinum* infected animals when treatment as initiated after granuloma colonization (Figure 2F).

### Opportunistic pathogens passively colonize primary *M. marinum* granulomas in zebrafish embryos

We examined the role of secondary mycobacterial infection in directing the colonization of primary granulomas by comparing the *M. abscessus* parental “low virulence” smooth colony morphotype to the more virulent rough colony morphotype used in our previous experiments. There was no difference in colonization by either colony morphotype compared to each other or relative to the lower rate of colonization seen in *M. marinum* superinfection (Supplementary Figure 2A). Interestingly, the rate of granuloma colonization was similar between WT and ΔESX1 *M. marinum* secondary infections, suggesting the differentiating factor between *M. marinum* and *M. abscessus* granuloma colonization potential is species-rather than virulence factor-specific. Furthermore, we did not observe a growth advantage for secondary *M. marinum* infection compared to infection into naïve embryos (Figure 3A).

**Figure 3.**
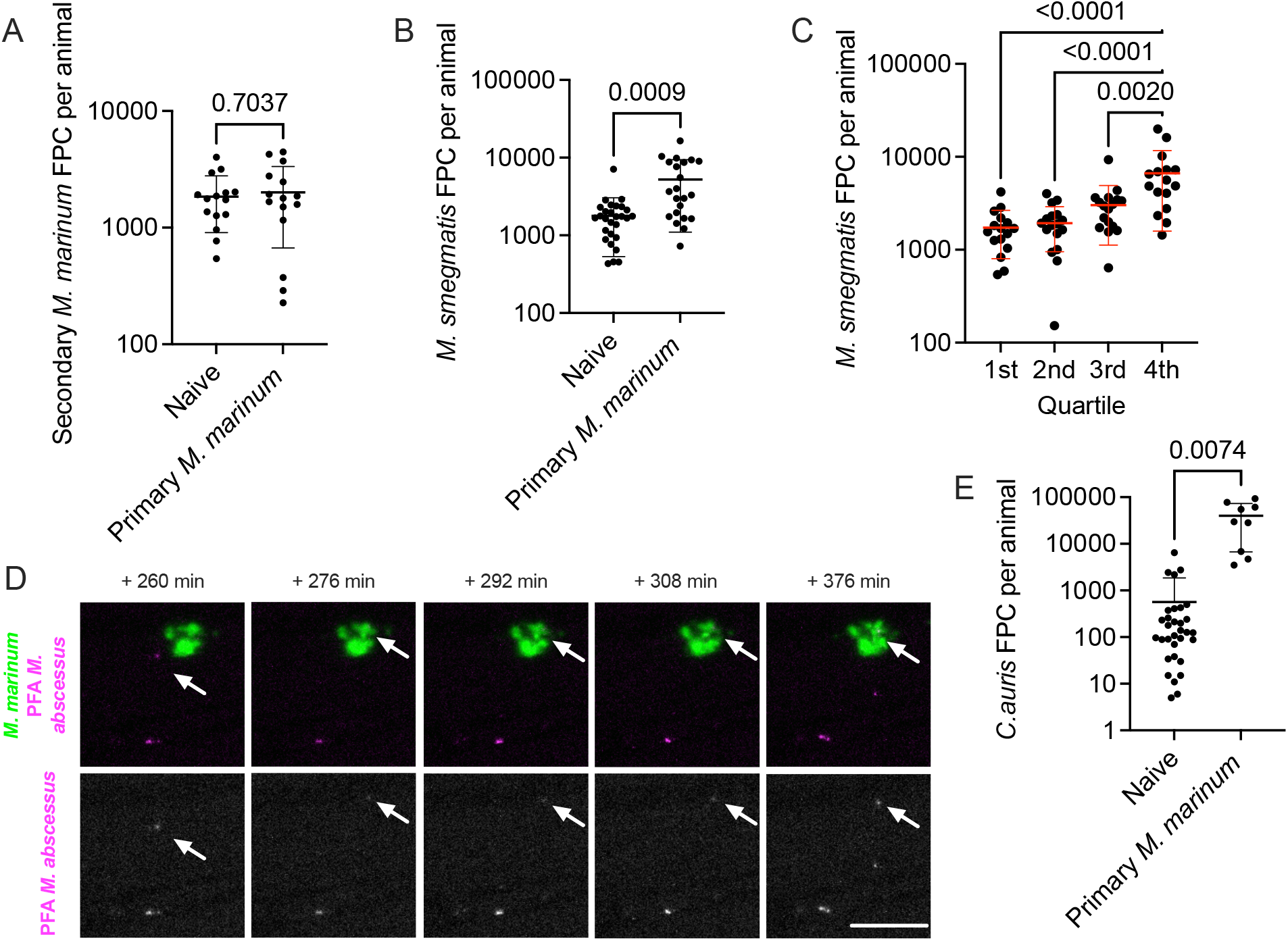
Colonization of primary granulomas provides a growth advantage to opportunistic pathogens A.Quantification of secondary *M. marinum* burden in 3 dpsi embryos infected with primary *M. marinum*. B.Quantification of secondary *M. smegmatis* burden in 3 dpsi embryos infected with primary *M. marinum*. C. Quantification of secondary *M. smegmatis* burden in 3 dpsi embryos infected with primary *M. marinum* grouped by quartile of *M. smegmatis* found within *M. marinum* granulomas. D. Images extracted from Supplementary Video 2 tracking ingress of PFA-fixed *M. abscessus* into a primary *M. marinum* granuloma from 260-276 minutes post tracking and residency of PFA-fixed *M. abscessus* inside granuloma for over duration of video. Arrow indicates location of PFA-fixed *M. abscessus* fluorescent signal, scale bar represents 100 μm. E. Quantification of secondary *C. auris* burden in 3 dpsi embryos infected with primary *M. marinum*.

We next performed super infection with fluorescent *Mycobacterium smegmatis*, typically considered avirulent. Similar to *M. abscessus*, superinfecting *M. smegmatis* had a significant growth advantage compared to infection into age-matched naïve embryos (Figure 3B), and total *M. smegmatis* burden was higher in the embryos in the highest quartile of granuloma colonization compared to the other three quartiles (Figure 3C). In co-injection studies, we observed a similar lack of protective effect of *M. marinum* on *M. smegmatis* burden in the absence of pre-existing granulomas and an antagonistic effect of *M. smegmatis* on *M. marinum* burden (Supplementary Figure 1B and 1C).

We also injected embryos with paraformaldehyde-killed *M. abscessus* and observed colonization of granulomas, suggesting the delivery of NTM to primary tuberculous granulomas is a passive feature of NTM species (Figure 3D, Supplementary Video 2).

This led us to test the bounds of the colonization phenotype by non-mycobacterial secondary infections. First, we used uropathogenic *Escherichia coli* (UPEC) as the secondary infection. This typically acute infection is capable of persisting in zebrafish embryos but we did not observe cross-genus interaction, consistent with data from a natural pathogen *Salmonella* experiment suggesting a different niche from NTMs (*8*) (Supplementary Figure 2B). Next, as fungal infections are a common consequence of post-TB lung disease and opportunistic yeast have the ability to survive within macrophages (*24*), we tested the ability of the *Candida auris* to colonize granulomas. We observed *C. auris* colonization of primary *M. marinum* granulomas and a growth advantage for *C. auris* in animals with an existing *M. marinum* infection (Figure 3E, Supplementary Figure 2C).

Together, these data suggest that granuloma colonization depends on the ability of secondary infections to be recognized by macrophages, survive intracellular killing mechanisms, but avoid overstimulating the macrophage to stop and form aggregates.

### ESX-1 directed maturation of primary *M. marinum* granulomas permits growth of superinfecting *M. abscessus*

The reproduction of the caseating granulomas seen in TB is a key advantage of the zebrafish- *M. marinum* infection model, and granulomas formed following trunk injection of *M. marinum* into embryos undergo stereotypical progression from cellular to necrotic granuloma from 3-5 dppi (*17, 21*). Reducing the primary inoculum and comparing the start of secondary infection from 3 to 4 dppi revealed a granuloma maturation-dependent increase in primary granuloma colonization by superinfecting *M. abscessus* (Figure 4A).

**Figure 4.**
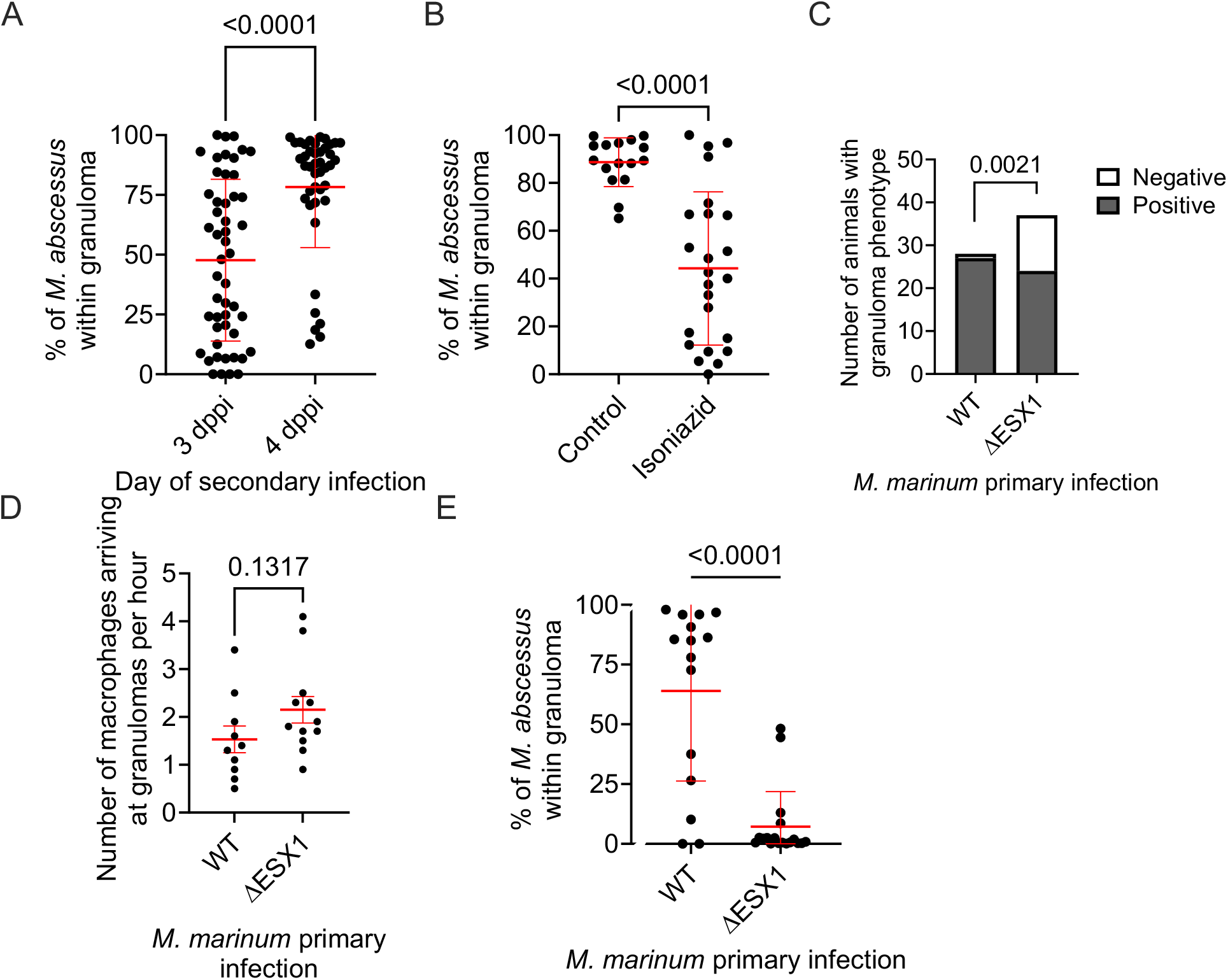
Secondary *M. abscessus* growth in primary granulomas is proportional to *M. marinum* ESX1-dependent granuloma maturation A.Quantification of *M. abscessus* colonization of primary *M. marinum* granulomas at 3 dpsi when infected at 3 or 4 dppi. B.Quantification of *M. abscessus* colonization of primary *M. marinum* granulomas at 3 dpsi treated with isoniazid. C.Quantification of animals with at least one *M. abscessus* colonization event of primary WT and ΔESX1 *M. marinum* granulomas at 1 dpsi. Data is pooled from two biological replicates. Statistical analysis reported is a Fisher’s Exact Test. D.Quantification of macrophage recruitment to WT vs ΔESX1 *M. marinum* granulomas across 4-22 hpsi. E.Quantification of *M. abscessus* colonization of primary WT and ΔESX1 *M. marinum* granulomas at 3 dpsi.

Conversely, arresting the maturation of primary granulomas with *M. marinum-*bacteriostatic isoniazid treatment at the time of secondary infection reduced the rate of primary granuloma colonization by superinfecting *M. abscessus* (Figure 4B).

The ESX1 type VII secretion system is the most important pro-granulomatous virulence factor in *M. marinum* and *M. tuberculosis* responsible for directly subverting phagocyte function, and driving necrosis and the recruitment of permissive macrophages to feed the granuloma (*25*). We found impaired colonization of primary ΔESX1 *M. marinum* granulomas compared to primary WT *M. marinum* granulomas early in infection even when similar burdens are reached with a higher initial inoculum (Figure 4C). Interestingly, we observed similar rate of macrophage recruitment to 5 dppi primary ΔESX1 *M. marinum* granulomas which contrasts to the reduced rate of macrophage recruitment early in ΔESX1 *M. marinum* infection compared to WT *M. marinum* infection (*26*), following secondary *M. abscessus* infection suggesting reduced macrophage migration is not the only factor accounting for reduced colonization of primary ΔESX1 *M. marinum* granulomas by secondary *M. abscessus* (Figure 4D). The small difference in initial colonization rate at 1 dpsi was compounded during later stages of *M. abscessus* superinfection with primary ΔESX1 *M. marinum* granulomas carrying only a very small minority of *M. abscessus* by 3 dpsi suggesting a requirement for ESX1-mediated necrosis in creating primary granulomas that attract and then further support the growth of superinfecting *M. abscessus* (Figure 4E)

We next investigated the mode of *M. abscessus* ingress into primary *M. marinum* granulomas by live imaging. As our sequential infection system introduced *M. abscessus* into the circulation, we first tracked the vascular source of *M. abscessus* ingression into *M. marinum* granulomas in the *Tg(kdrl:egfp)*^*s843*^ line, where vascular endothelial cells are labelled by EGFP expression (*27*). We observed extravasation of *M. abscessus* from blood vessels adjacent and distal to the granuloma followed by directional migration and ingress (Supplementary Video 1). Ingress events originating from adjacent intersegmental and dorsal longitudinal anastomotic vessels were classed as local vasculature and all other vessels were classed as distal vasculature (Supplementary Figure 2A). Quantification of 114 extravasation events leading to *M. abscessus* ingress in 11animals revealed a mix of 60 local and 54 distal extravasation events, demonstrating the possibility of both hematogenous spread and macrophage carriage.

We have previously used vascular normalization as a host-directed therapy to reduce the extravasation of neutrophils around *M. marinum* granulomas. Here, we hypothesized that preventing vascular pathology would reduce the extravasation of *M. abscessus* towards granulomas in our superinfection model (*28*). We treated *M. marinum*-infected embryos with pazopanib, an FDA-approved VEGFR inhibitor with host-directed activity against *M. marinum* infection-induced vascular pathologies (*21*), starting one day prior to secondary *M. abscessus* infection and continuing until 3 dpsi. Treatment with pazopanib reduced the rate of primary granuloma colonization by superinfecting *M. abscessus* when assayed 3 dpsi (Supplementary Figure 2B).

Previous live imaging and histological studies have implicated ingress of infected macrophages as the primary mode of *M. abscessus* and *M. marinum* dissemination in zebrafish. Our extravasation imaging demonstrated the direct migration of *M. abscessus* into granulomas, suggesting macrophage carriage (*29-32*). To confirm this hypothesis, we performed live imaging of granuloma ingress in the *Tg(acod1:tdtomato)*^*xt40*^ line, where macrophages are labelled with TdTomato (Supplementary Video 3), and carriage of *M. abscessus* by GFP positive and negative cells with similar DIC morphologies in the alternative *TgBAC(mpeg1*.*1:egfp)*^*vcc7*^ where a subset of macrophages are labelled with EGFP (Supplementary Video 4) (*33*).

Neutrophils are abundant immune cells that play a crucial role in controlling *M. abscessus* and *M. marinum* in zebrafish embryos (*32, 34, 35*). To determine the potential contribution of neutrophil carriage, we performed live imaging of granuloma ingress in the *Tg(lyzc:egfp)*^*nz117*^ line, where neutrophils are labelled with EGFP (*36*). As expected, we observed neutrophil recruitment to primary granulomas and interactions with *M. abscessus*. However, GFP-positive neutrophil carriage of *M. abscessus* was rare (Supplementary Video 5). Quantification of 75 *M. abscessus* ingress events in 15 animals revealed only 6 events, demonstrating a low utilization of neutrophils for carriage of *M. abscessus*.

### *M. abscessus* adaptation to primary granuloma residency

To understand how *M. abscessus* takes advantage of the necrotic granuloma niche we first examined the transcriptome of *M. abscessus* grown in complete 7H9 media with *M. abscessus* in the zebrafish embryo infection model. Examination of known stress response gene families: Esx-3, mycobactins, oxidative stress, and cell wall remodeling confirmed the zebrafish embryo infection model induces the expression of a virulence-associated *M. abscessus* gene program (Figure 5A). Further comparison of the *M. abscessus* lipid metabolism gene expression program revealed an upregulation of lipid metabolism during adaptation to infection suggesting *M. abscessus* may utilize the lipid-rich caseum to fuel accelerated growth during superinfection (Figure 5B).

**Figure 5.**
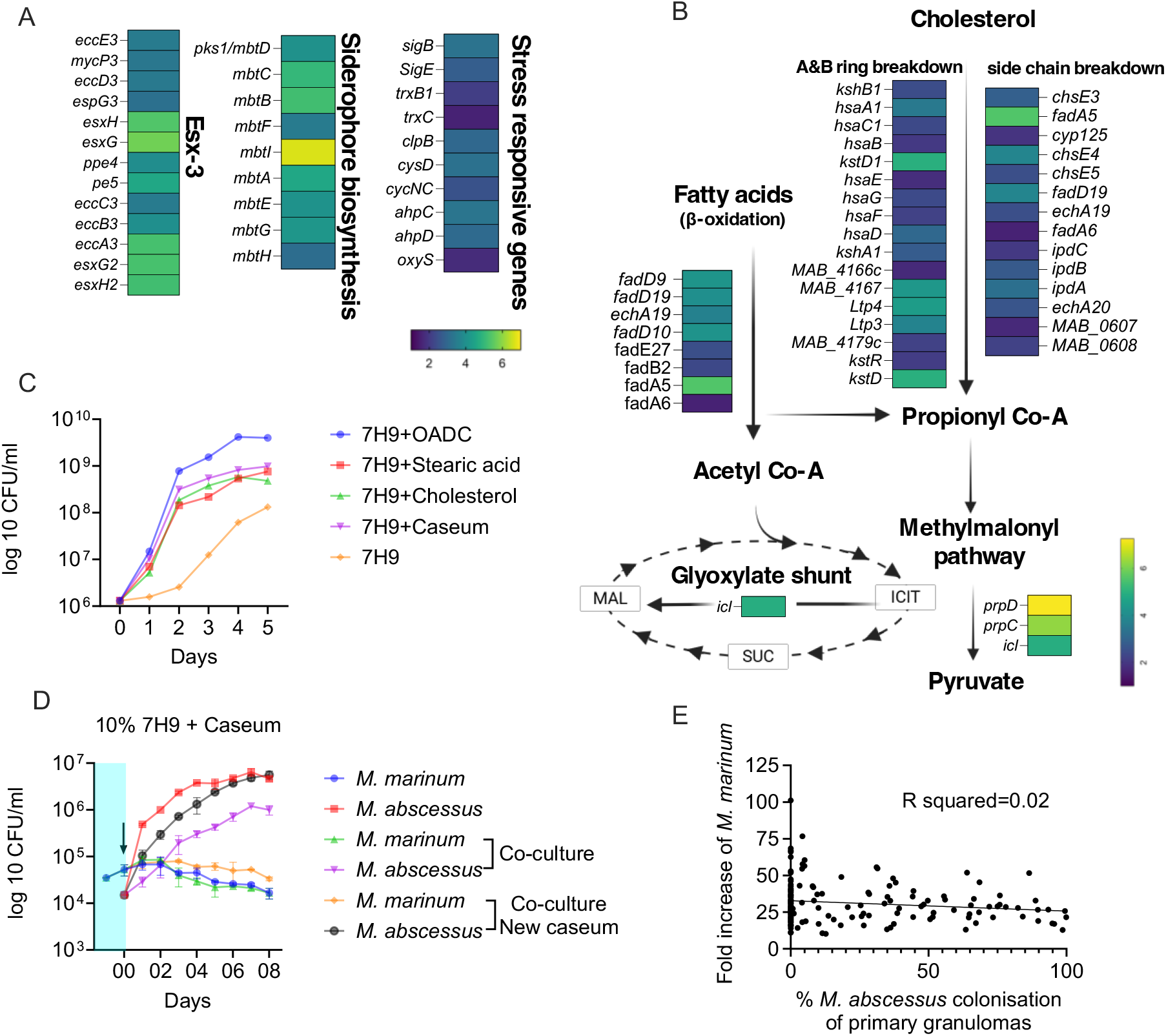
*M. abscessus* occupies a separate niche to *M. marinum* within the necrotic granuloma A.Heat map of *M. abscessus* stress gene expression derived from RNAseq of *M. abscessus* at 6 dpi in zebrafish embryos compared to baseline *in vitro* 7H9 broth cultured *M. abscessus*. B.Heat map of *in vivo M. abscessus* lipid metabolism gene expression compared to *in vitro* 7H9 broth cultured *M. abscessus*. C.Quantification of *M. abscessus* growth in 7H9 media supplemented with carbon sources. D.Quantification of *M. abscessus* and *M. marinum* growth in 7H9 media diluted to 10% with PBS and supplemented with *in vitro* caseum. *M. marinum* culture was performed for 4 days prior to Day 00 when *M. abscessus* is added to the co-culture conditions. E.Quantification of *M. marinum* growth in 163 individual embryos from 0 to 3 dpsi with *M. abscessus*. R squared value calculated by linear regression, slope equation Y = -0.07066*X + 32.79, P=0.0481.

To test the hypothesis that *M. abscessus* could adapt to the caseum in the necrotic core as a nutrient source, we next switched to an *in vitro* system. We analyzed the growth of *M. abscessus* in diluted 7H9 media supplemented with OADC as a positive control, RAW 264.7 cell-derived *in vitro* caseum (*37*), and the individual lipid substrates cholesterol and stearic acid. Growth curves demonstrated that *M. abscessus* is able to grow rapidly with a range of lipid substrates, specifically being able to utilize *in vitro* caseum as a substrate (Figure 5C).

To model our *in vivo* sequential infections, we first cultured *M. marinum* in *in vitro* caseum, facilitating pre-conditioning of the caseum. After 4 days when *M. marinum* growth had plateaued, we introduced *M. abscessus* to the culture system. The addition of *M. abscessus* to the established *M. marinum* culture resulted in lower overall growth of *M. abscessus* compared to its monoculture in sterile caseum consistent with the depletion of some nutrients by *M. marinum* pre-conditioning (Figure 5D). This growth defect was ameliorated when new caseum was added to the established *M. marinum* culture system, suggesting that the optimal *M. abscessus* growth conditions overlap with nutrients utilized by *M. marinum* (Figure 5D). We observed similar stability of *M. marinum* CFU levels across all experiments, regardless of whether *M. abscessus* was added to the culture, suggesting a neutral interpretation of interaction between the two species by *M. marinum*.

To study this interaction *in vivo*, we analyzed the growth of *M. marinum* in individual embryos with and without sequential *M. abscessus* infection. Analysis of embryos infected with only *M. marinum* revealed a wide range (11-101x) of *M. marinum* growth across days 3-6 post primary infection, equivalent to 0-3 dpsi in the sequential infection experiment, and all sequential infection values fell within this range (Figure 5E). Further analysis of *M. marinum* fold change from 0-3 dpsi infection demonstrated no correlation between *M. marinum* growth and the rate of *M. abscessus* colonization of primary *M. marinum* granulomas (Figure 5E). Together, these data demonstrate that *M. abscessus* occupies a unique niche in necrotic granulomas that is not at the expense of *M. marinum*.

## Discussion

In Singapore, a country that has effectively eliminated local transmission but retains the epidemiological “scar” of historically high prevalence of TB, *M. abscessus* ais one of the most prevalent NTM infections, with prior TB being the most common predisposing factor (*38*). Evidence from mammalian models suggests that increasing TB severity drives a shift from leukocyte-mediated heterologous protection against secondary mycobacterial infection afforded by a contained primary infection toward systemic reprogramming of hematopoiesis, leading to increased susceptibility to secondary mycobacterial infection following disseminated disease (*14, 15, 39*). These observations suggest that compromised leukocyte immunity could drive increased NTM susceptibility in treated TB patients

### Tuberculous granulomas are a predisposing factor for opportunistic NTM infection

Our data provide compelling evidence that granulomas formed during TB infection provide a niche for the growth of *M. abscessus* in otherwise immunologically intact resistant hosts. This finding has significant implications for explaining the increased risk of NTM infection in patients with prior or ongoing pulmonary TB. Post TB lung disease encompasses a wide range of histological lung remodeling, leading to a loss of respiratory function, which is hypothesized to impair the physical clearance of NTM following inoculation from environmental reservoirs (*3*). Although our models are unable to replicate the physical clearance of inhaled NTM, they faithfully produce TB-like granulomas with caseous necrosis. We posit that unresolved TB granulomas should be considered a susceptibility factor for infection by opportunistic pathogens.

The rapid growth of *M. abscessus* and *M. marinum* in co-culture experiments are most likely due to the relative growth rates of the organisms rather than a true virulence advantage. Dual-species granulomas are usually bounded by *M. marinum* or exhibit a mixed fluorescent signal, suggesting *M. abscessus* does not expand these histopathological structures in our assays. Furthermore, naïve embryos were largely able to control *M. abscessus* when infected at 5 dpf but had early granuloma formation when infected with *M. marinum*, consistent with *M. marinum* having a much higher relative growth potential per unit of inoculum across zebrafish infection assays (*16, 21*). Further experiments in appropriate model systems will be required to determine the effect of primary tuberculous infection clearance from granulomas on the survival of super infecting NTM who can no longer “hide” behind ESX1 immune subversion driven by the primary species.

Our findings highlight the importance of efforts to find preventative and interventional treatments for TB pathologies, including granuloma resolution, as the risks of post-infection sequalae will persist long after TB transmission is eradicated. Many NTM cases are initially diagnosed as TB, resulting in empirical anti-TB therapy to which *M. abscessus* is insensitive. These cases can be variably defined as recent, prior, or concomitant TB with *M. abscessus* infection, depending on the level of specificity in the initial diagnosis. In cases where there is evidence of *M. tuberculosis* at the first diagnosis, our findings suggest that these patients may be at risk of severe *M. abscessus* infection phenotypes, such as fibrocavitary disease compared to the bronchial nodular form.

### When does primary TB increase resistance to secondary mycobacterial infection?

While there is mixed evidence from preclinical infection models for and against a protective effect of primary *M. tuberculosis* infection on reinfection, epidemiological evidence clearly demonstrates that prior *M. tuberculosis* or NTM infection is a risk factor for future tuberculous and non-tuberculous mycobacterial infection (*38, 40, 41*). While some of this risk can be attributed to patients remaining in physical environments with high mycobacterial exposure levels, countries such as Singapore, where TB transmission was effectively eliminated within a generation, manifest a long, lingering tail of post-TB susceptibility to NTM infection at a population level (*38, 42*).

The severity of primary infection in preclinical models appears to be the decisive factor for determining if primary mycobacterial infection is protective against future challenge. Naturally controlled murine and non-human primate *M. tuberculosis* infections clearly protect against sequential *M. tuberculosis* infection (*15, 18*). Poor initial control of infection by highly virulent strains, systemic administration of live *M. tuberculosis* (*39, 43*), or the converse clearing of infection (*14, 15*) can compromise immunological control of the second infection by a range of mechanisms from the hematopoietic stem cell through to the pulmonary microenvironment. This suggests a Goldilocks principle whereby ongoing inflammatory and antigenic stimulation is the primary driver of protective anti-mycobacterial immune responses. It will be important to adapt these models to investigate the relative contributions of hematopoietic reprogramming and pulmonary adaptive immunity to the restriction of NTM in models that recapitulate human-like granulomas.

## Methods

### Zebrafish

Zebrafish experiments were carried out under the A*STAR IACUC approvals 211667 and 221694. Adults were housed under 14 hour light / 10 hour dark cycles in 28°C recirculating systems. Embryos were produced by natural spawning and raised in E3 media supplemented with PTU at 28°C.

### Growth of microbes

*M. abscessus, M. marinum*, and *M. smegmatis* were cultured in 7H9 or on 7H10 supplemented with OADC and hygromycin to select for pTEC fluorescent protein plasmids (L. Ramakrishnan, plasmids are available through Addgene https://www.addgene.org/Lalita_Ramakrishnan/). *C. auris* expressing mCherry was prepared as previously described and is available upon request from EWLC (*44*). Uropathogenic *E. coli* was prepared as previously described (*45*).

PFA killing of *M. abscessus* was carried out by resuspending a midlog culture of *M. abscessus* in 4% PFA in PBS for 30 minutes at room temperature. Fixed bacteria were rinsed prior to injection. Validation of killing was performed by plating on 7H10 supplemented with OADC and incubation at 37°C for 7 days.

Growth curves of *M. abscessus* and *M. marinum* were performed in 10% 7H9 media diluted with PBS at 30°C in a static incubator. *In vitro* caseum was produced as previously described (*37*).

### Adult zebrafish infections

Adult zebrafish were injected with approximately 100 CFU *M. marinum* or 5×10^5^ to 10^6^ CFU of *M. abscessus* as indicated. Animals were housed under 14 hour light / 10 hour dark cycles in an isolated 28°C recirculating system and fed once daily with a nutritionally complete dry feed.

Bacterial enumeration by CFU recovery was performed by bead beating individual adult zebrafish in a Tomy Micro Smash MS-100 and plating of homogenate on 7H10 for *M. marinum* or LB Agar for *M. abscessus* supplemented with hygromycin. Growth of *M. marinum* was carried out at 30°C for 7 days and growth of *M. abscessus* was carried out at 37°C for 5 days.

Histology was carried out using a Leica CM1520 to cut 10 μm thick cryosections of PFA-fixed adults and counterstaining with DAPI. Fluorescent images of preserved bacterial fluorescence were acquired using a Nikon Ni-E upright microscope.

### Zebrafish embryo infections

Zebrafish embryos were infected by microinjection with approximately 200 CFU *M. marinum*, 1000 CFU ∧1ESX1 *M. marinum* or *M. abscessus*. Primary infection was performed by injection into the neural tube above the yolk sac extension at 2 dpf and secondary infection was performed by intravascular injection into the dorsal aorta or caudal vein at 5 dpf/3 dpi, unless otherwise described.

Clodronate microsomes were co-injected with secondary *M. abscessus* infection by intravascular injection into the dorsal aorta or caudal vein at 5 dpf/3 dpi.

Clarithromycin (10 μg/ml final concentration), isoniazid (50 μM final concentration), pazopanib (500 nM final concentration) were added directly to embryo media.

### Microscopy of zebrafish embryos

All microscopy was carried out stereomicroscopy of immobilized zebrafish embryos. Static imaging was performed on a Nikon SMZ25 stereoscope or an inverted Nikon Eclipse Ti2. Time lapse imaging was performed on inverted Thermofisher EVOS7000 or Olympus IX-83 microscopes.

Image analysis was performed in ImageJ/FIJI with bacterial fluorescent pixel count (FPC)carried out as previously described (*46*). Percentage of *M. abscessus* in granulomas was calculated by the formula (*M. abscessus* FPC overlap with *M. marinum* / total *M. abscessus* FPC) x 100.

### RNA sequencing of *M. abscessus*

Mycobacterial RNA was harvested from 3 mid-log 7H9 cultures and 3 pools of trizol-pre lysed 7 dpi *M. abscessus-*infected zebrafish embryos by bead beating in Lysis buffer and further processed according to manufactures protocol (MN-NucleoSpin RNA kit). Total RNA was subjected to 150 bp paired end sequencing on an Illumina Novaseq 6000 (NovogeneAIT Genomics Singapore). Sequencing quality check was carried out by FASTQC. Alignment of read pairs to *Mycobacteroides abscessus* genome (ASM6918v1, GenBank) was performed by STAR. The raw read count matrix was generated by featureCounts. DESeq2 analysis was performed to identify differentially expressed genes (DEGs). DEGs were manually annotated to stress response and lipid metabolism pathways for display.

### Statistics

All analyses of infection experiments were carried out with Graphpad Prism using T-tests for pairwise comparisons or ANOVA for multiple comparisons. All data are representative of at least 3 biological replicates unless otherwise stated in the captions. Error bars on graphs represent standard deviation.

## Acknowledgements

This study was funded by the Singapore Ministry of Health’s National Medical Research Council under its individual research grant scheme (OFIRG22jul-0081) to S.H.O.

A*STAR IMCB Aquarium Platform for expert zebrafish husbandry. Dr Eloise Ma and the A*STAR Microscopy Platform for microscopy assistance.

Dr J Muse Davis for helpful discussion of live imaging techniques. Professors Lalita Ramakrishnan and Paul Edelstein for discussion of co-infection and sequential infections. Members of A*STAR ID Labs and the SG BUG community for discussion.

---

Supplementary Video 1

Time lapse imaging of magenta *M. abscessus* ingress into a yellow *M. marinum* granuloma in a 5 dppi *Tg(kdrl:egfp)* zebrafish larva with green blood vessels.

Supplementary Video 2

Time lapse imaging of magenta PFA-fixed *M. abscessus* ingress into a green *M. marinum* granuloma in a 5 dppi zebrafish larva.

Supplementary Video 3

Time lapse imaging of cyan *M. abscessus* ingress into a yellow *M. marinum* granuloma in a 5 dppi *Tg(acod1:tdtomato)*^*xt40*^ zebrafish larva with red macrophages.

Supplementary Video 4

Time lapse DIC imaging merged with fluorescent imaging of magenta *M. abscessus* being carried by green macrophages in a 5 dppi *Tg(mpeg1*.*1:egfp)* zebrafish larva with cyan *M. marinum* granulomas.

Supplementary Video 5

Time lapse imaging of magenta *M. abscessus* ingress into a blue *M. marinum* granuloma in a 5 dppi *Tg(lyzC:gfp)* zebrafish larva with green neutrophils.

**Supplementary Figure 1.**
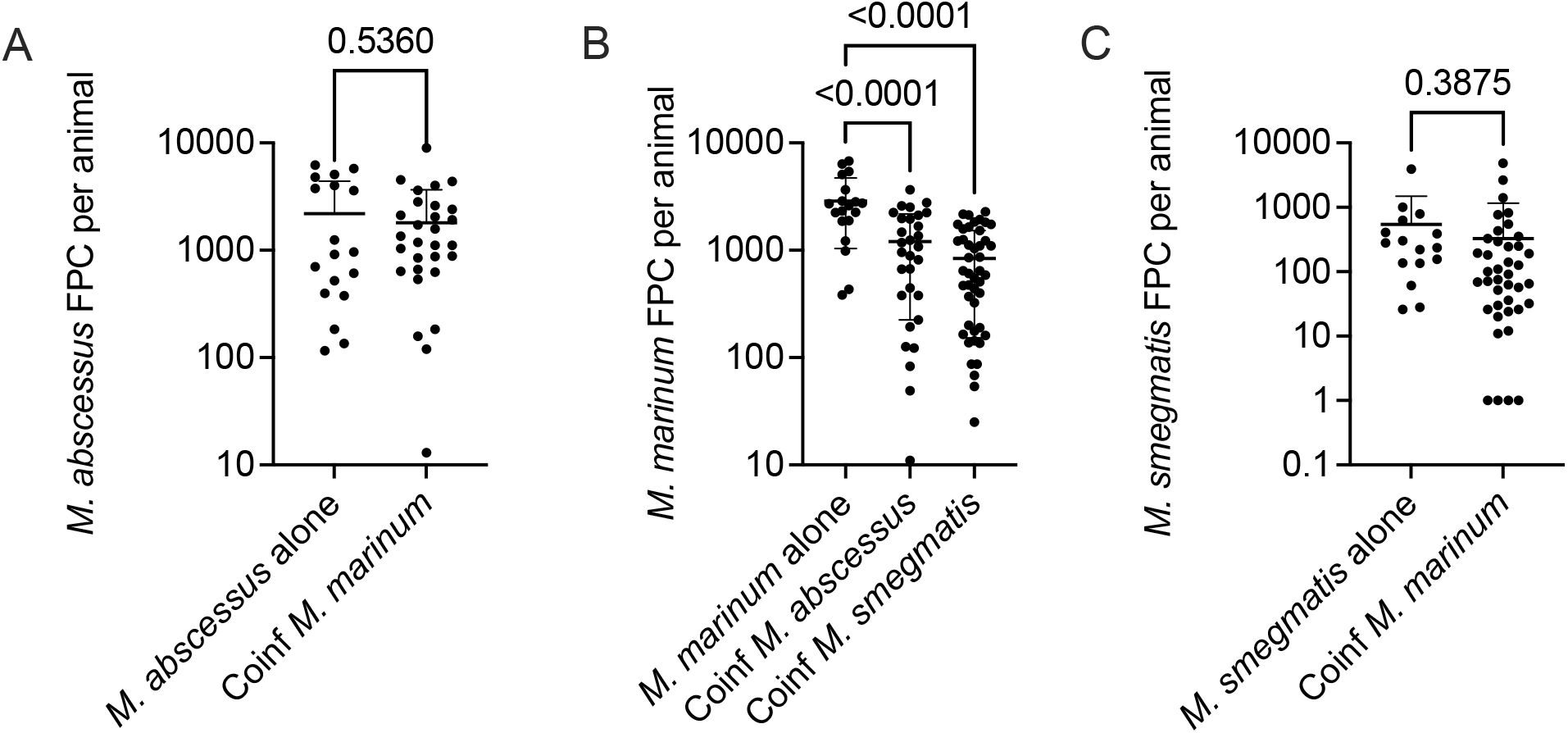
A. Quantification of *M. abscessus* burden in 4 days post infection embryos co-injected with *M. abscessus* and *M. marinum*. B. Quantification of *M. marinum* burden in 4 days post infection embryos co-injected with *M. abscessus* or *M. smegmatis* and *M. marinum*. C. Quantification of *M. smegmatis* burden in 4 days post infection embryos co-injected with *M. smegmatis* and *M. marinum*.

**Supplementary Figure 2.**
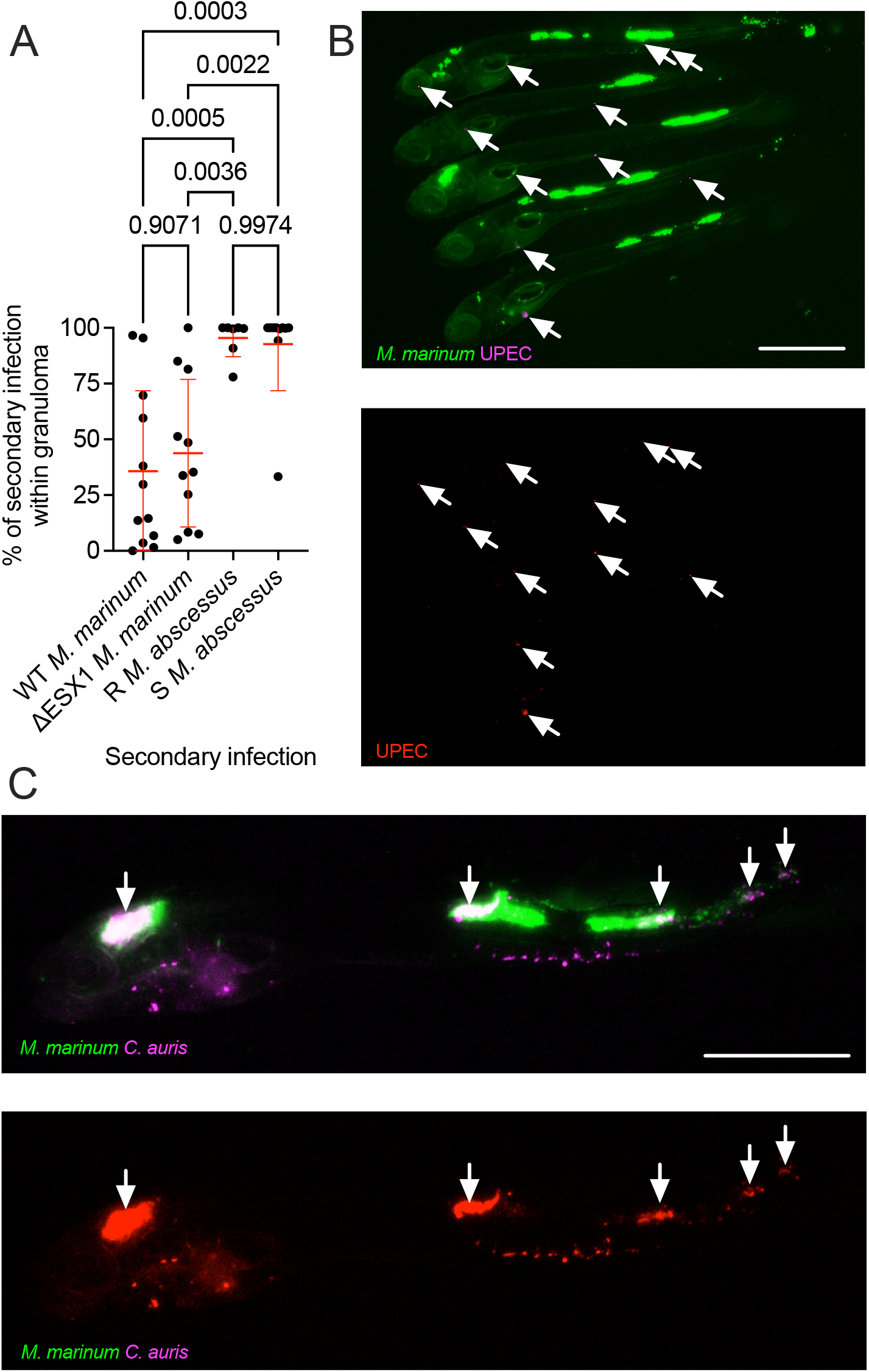
A.Quantification of *M* .*marinum* WT, ΔESX1 *M. marinum*, R and S *M. abscessus* colonization of primary *M. marinum* granulomas at 3 dpsi. B.Representative image of secondary infection UPEC distribution in zebrafish embryos with primary *M. marinum* infection at 3 dpsi. Arrows indicate locations of UPEC fluorescent signals, scale bar represents 100 μm. C.Representative image of secondary infection *C. auris* distribution in a zebrafish embryo with primary *M. marinum* infection at 3 dpsi. Arrows indicate locations of *C. auris* fluorescent signals within *M. marinum* granulomas, scale bar represents 100 μm.

**Supplementary Figure 3.**
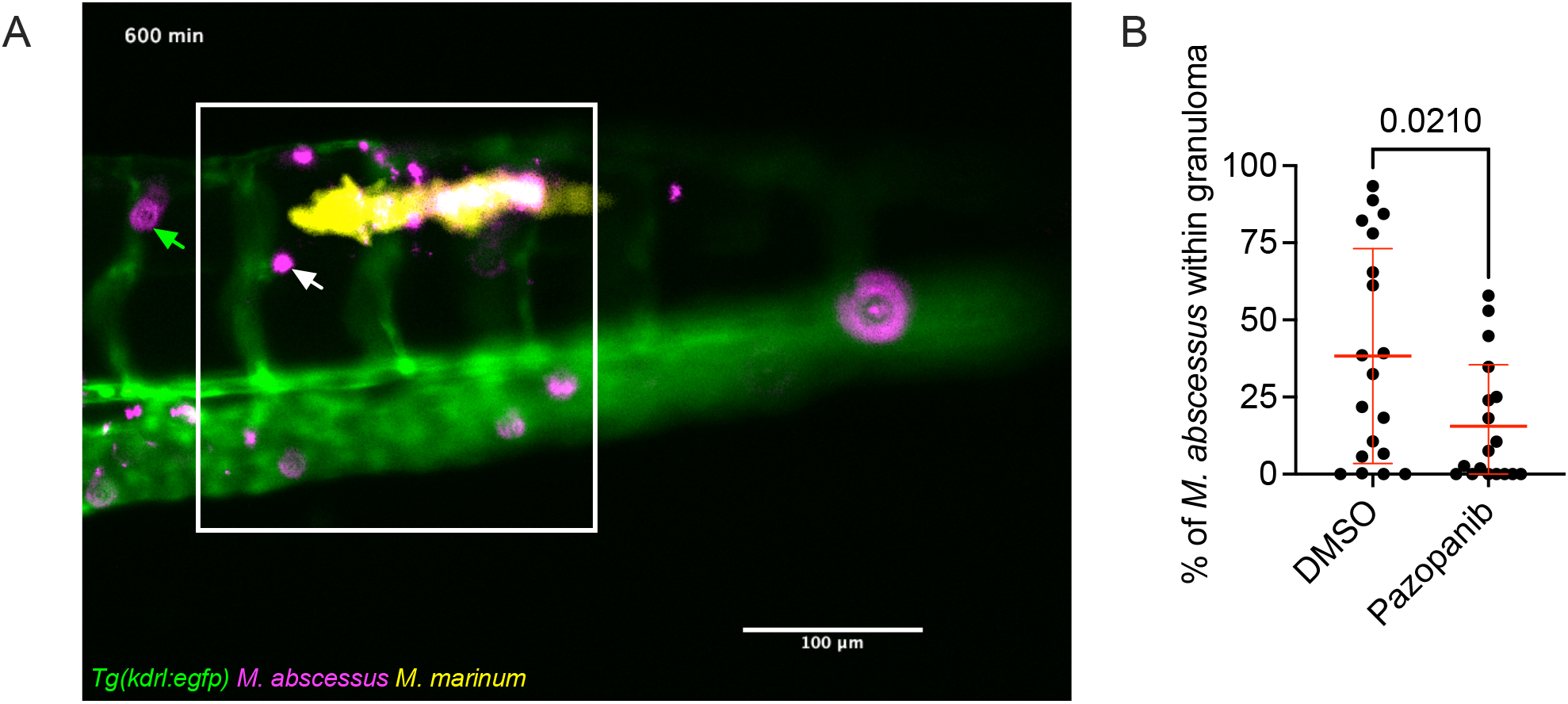
A.Still image extracted from Supplementary Video 1. White box demarcates area of local *Tg(kdrl:egfp)-*positive vasculature (green) around primary *M. marinum* granuloma (yellow). Extravasating *M. abscessus* (purple) are indicated by white arrow for local vasculature or green arrow for distal vasculature. B.Quantification of *M. abscessus* colonization of primary *M. marinum* granulomas at 3 dpsi treated with pazopanib.

## References

1. M. D. Johansen, J. L. Herrmann, L. Kremer, Non-tuberculous mycobacteria and the rise of Mycobacterium abscessus. Nat Rev Microbiol 18, 392–407 (2020).

2. G. J. Fox et al., Post-treatment Mortality Among Patients With Tuberculosis: A Prospective Cohort Study of 10 964 Patients in Vietnam. Clin Infect Dis 68, 1359–1366 (2019).

3. B. W. Allwood et al., Post-Tuberculosis Lung Disease: Clinical Review of an Under-Recognised Global Challenge. Respiration; international review of thoracic diseases 100, 751–763 (2021).

4. Y. He, J. L. Wang, Y. A. Zhang, M. S. Wang, Prevalence of Culture-Confirmed Tuberculosis Among Patients with Nontuberculous Mycobacterial Disease. Infect Drug Resist 15, 3097–3101 (2022).

5. C. K. Lin et al., Incidence of nontuberculous mycobacterial disease and coinfection with tuberculosis in a tuberculosis-endemic region: A population-based retrospective cohort study. Medicine (Baltimore) 99, e23775 (2020).

6. M. Silva Miranda, A. Breiman, S. Allain, F. Deknuydt, F. Altare, The tuberculous granuloma: an unsuccessful host defence mechanism providing a safety shelter for the bacteria? Clinical & developmental immunology 2012, 139127 (2012).

7. M. A. Behr, P. H. Edelstein, L. Ramakrishnan, Is Mycobacterium tuberculosis infection life long? BMJ 367, l5770 (2019).

8. C. L. Cosma, O. Humbert, L. Ramakrishnan, Superinfecting mycobacteria home to established tuberculous granulomas. Nat Immunol 5, 828–835 (2004).

9. C. L. Cosma, O. Humbert, D. R. Sherman, L. Ramakrishnan, Tracicking of superinfecting Mycobacterium organisms into established granulomas occurs in mammals and is independent of the Erp and ESX-1 mycobacterial virulence loci. J Infect Dis 198, 1851–1855 (2008).

10. M. Moreno-Molina et al., Genomic analyses of Mycobacterium tuberculosis from human lung resections reveal a high frequency of polyclonal infections. Nat Commun 12, 2716 (2021).

11. T. D. Lieberman et al., Genomic diversity in autopsy samples reveals within-host dissemination of HIV-associated Mycobacterium tuberculosis. Nat Med 22, 1470–1474 (2016).

12. H. Liu et al., Retrospective clinical and microbiologic analysis of metagenomic next-generation sequencing in the microbiological diagnosis of cutaneous infectious granulomas. Ann Clin Microbiol Antimicrob 23, 84 (2024).

13. W. Du et al., Association of bacteriomes with drug susceptibility in lesions of pulmonary tuberculosis patients. Heliyon 10, e37583 (2024).

14. S. K. Ganchua et al., Antibiotic treatment modestly reduces protection against Mycobacterium tuberculosis reinfection in macaques. Infect Immun, e0053523 (2024).

15. J. Nemeth et al., Contained Mycobacterium tuberculosis infection induces concomitant and heterologous protection. PLoS Pathog 16, e1008655 (2020).

16. J. Kam et al., Rough and smooth variant Mycobacterium abscessus infections are differentially controlled by host immunity during chronic infection of adult zebrafish. Nat Commun 13, 952 (2022).

17. L. E. Swaim et al., Mycobacterium marinum infection of adult zebrafish causes caseating granulomatous tuberculosis and is moderated by adaptive immunity. Infect Immun 74, 6108–6117 (2006).

18. J. D. Bromley et al., CD4(+) T cells re-wire granuloma cellularity and regulatory networks to promote immunomodulation following Mtb reinfection. Immunity, (2024).

19. M. R. Cronan et al., Macrophage Epithelial Reprogramming Underlies Mycobacterial Granuloma Formation and Promotes Infection. Immunity 45, 861–876 (2016).

20. M. R. Cronan et al., A non-canonical type 2 immune response coordinates tuberculous granuloma formation and epithelialization. Cell 184, 1757–1774 e1714 (2021).

21. S. H. Oehlers et al., Interception of host angiogenic signalling limits mycobacterial growth. Nature 517, 612–615 (2015).

22. J. P. Sarathy et al., A Novel Tool to Identify Bactericidal Compounds against Vulnerable Targets in Drug-Tolerant M. tuberculosis found in Caseum. mBio 14, e0059823 (2023).

23. M. Xie et al., ADP-ribosylation-resistant rifabutin analogs show improved bactericidal activity against drug-tolerant M. abscessus in caseum surrogate. Antimicrob Agents Chemother 67, e0038123 (2023).

24. J. L. Tenor, S. H. Oehlers, J. L. Yang, D. M. Tobin, J. R. Perfect, Live Imaging of Host-Parasite Interactions in a Zebrafish Infection Model Reveals Cryptococcal Determinants of Virulence and Central Nervous System Invasion. mBio 6, (2015).

25. C. L. Cosma, K. Klein, R. Kim, D. Beery, L. Ramakrishnan, Mycobacterium marinum Erp is a virulence determinant required for cell wall integrity and intracellular survival. Infect Immun 74, 3125–3133 (2006).

26. J. M. Davis, L. Ramakrishnan, The role of the granuloma in expansion and dissemination of early tuberculous infection. Cell 136, 37–49 (2009).

27. S. W. Jin, D. Beis, T. Mitchell, J. N. Chen, D. Y. Stainier, Cellular and molecular analyses of vascular tube and lumen formation in zebrafish. Development 132, 5199–5209 (2005).

28. J. Y. Kam et al., Inhibition of infection-induced vascular permeability modulates host leukocyte recruitment to Mycobacterium marinum granulomas in zebrafish. Pathogens and disease 80, (2022).

29. H. Clay et al., Dichotomous role of the macrophage in early Mycobacterium marinum infection of the zebrafish. Cell Host Microbe 2, 29–39 (2007).

30. J. M. Davis et al., Real-time visualization of mycobacterium-macrophage interactions leading to initiation of granuloma formation in zebrafish embryos. Immunity 17, 693–702 (2002).

31. A. Bernut et al., Mycobacterium abscessus cording prevents phagocytosis and promotes abscess formation. Proc Natl Acad Sci U S A 111, E943–952 (2014).

32. 1. A. Bernut et al., Mycobacterium abscessus-Induced Granuloma Formation Is Strictly Dependent on TNF Signaling and Neutrophil Tracicking. PLoS Pathog 12, e1005986 (2016).

33. W. J. Brewer et al., Macrophage NFATC2 mediates angiogenic signaling during mycobacterial infection. Cell reports 41, 111817 (2022).

34. P. M. Elks et al., Hypoxia Inducible Factor Signaling Modulates Susceptibility to Mycobacterial Infection via a Nitric Oxide Dependent Mechanism. PLoS Pathog 9, e1003789 (2013).

35. C. T. Yang et al., Neutrophils exert protection in the early tuberculous granuloma by oxidative killing of mycobacteria phagocytosed from infected macrophages. Cell Host Microbe 12, 301–312 (2012).

36. C. Hall, M. V. Flores, T. Storm, K. Crosier, P. Crosier, The zebrafish lysozyme C promoter drives myeloid-specific expression in transgenic fish. BMC Dev Biol 7, 42 (2007).

37. J. P. Sarathy et al., An In Vitro Caseum Binding Assay that Predicts Drug Penetration in Tuberculosis Lesions. J Vis Exp, (2017).

38. A. Y. H. Lim et al., Profiling non-tuberculous mycobacteria in an Asian setting: characteristics and clinical outcomes of hospitalized patients in Singapore. BMC pulmonary medicine 18, 85 (2018).

39. N. Khan et al., M. tuberculosis Reprograms Hematopoietic Stem Cells to Limit Myelopoiesis and Impair Trained Immunity. Cell 183, 752–770 e722 (2020).

40. X. Shen et al., Recurrent tuberculosis in an urban area in China: Relapse or exogenous reinfection? Tuberculosis (Edinb) 103, 97–104 (2017).

41. S. Verver et al., Rate of reinfection tuberculosis after successful treatment is higher than rate of new tuberculosis. Am J Respir Crit Care Med 171, 1430–1435 (2005).

42. Z. X. Zhang, B. P. Z. Cherng, L. H. Sng, Y. E. Tan, Clinical and microbiological characteristics of non-tuberculous mycobacteria diseases in Singapore with a focus on pulmonary disease, 2012-2016. BMC infectious diseases 19, 436 (2019).

43. M. Henao-Tamayo et al., A mouse model of tuberculosis reinfection. Tuberculosis (Edinb) 92, 211–217 (2012).

44. J. Gao et al., LncRNA DINOR is a virulence factor and global regulator of stress responses in Candida auris. Nat Microbiol 6, 842–851 (2021).

45. K. Wright et al., Mycobacterial infection-induced miR-206 inhibits protective neutrophil recruitment via the CXCL12/CXCR4 signalling axis. PLoS Pathog, (2021).

46. M. A. Matty, S. H. Oehlers, D. M. Tobin, Live Imaging of Host-Pathogen Interactions in Zebrafish Larvae. Methods Mol Biol 1451, 207–223 (2016).

